# The role of Vitamin C in the energy supply of cells Hypothetical structure for energy transformation

**DOI:** 10.1101/2020.07.21.214403

**Authors:** János Hunyady

**Affiliations:** Department of Dermatology, Faculty of Medicine, University of Debrecen., Nagyerdei krt. 98, H-4032, Debrecen, Hungary

**Keywords:** NAD, FAD, ascorbic acid, uric acid, glucose

## Abstract

The present paper represents a hypothetical structure, the structure for energy transformation (SET), which might be responsible for the proper energy transformation steps leading to the continuous production of H^+^ and ATP in living cells. We predict that the electron flow is realized through the electron flow device (EFD). We suppose that there are several versions of the SET. Two of them are described below [Structure of Aerobic Glycolysis (SET-AG) and Structure of Oxidative Phosphorylation (SET-OP)]. The hypothesis is based on the atom properties of the protonated adenine molecule and the docking computations of molecular mechanics involved, suggesting that two ascorbate molecules may occupy the empty NADPH pocket, preferably binding to the adenine binding site. We hypothesize that the adenine originates from uric acid (UA), resulting in an ATP-UA-ADP-ATP-UA cycle. It would also mean that UA is one of the oxygen sources in aerobic glycolysis.

We also suppose that the EFD contains the well-known molecules of the Nicotinamide Adenine Dinucleotide (NAD) and Flavin Adenine Dinucleotide (FAD) supplemented with two additional UA-originated adenine molecules, two L-ascorbic acids, and two D-glucose molecules. Based on all this, we surmise that a tetra adenine octo phosphate ring (TAR) exists, in which the UA originated adenine molecules form a ring. The molecules are linked to each other through the N7-C2 and C8-N1 atoms of the adenine molecules by H_2_PO_4_^e-^ molecules. The four N10 atoms of the adenine molecules bind one flavin, one nicotinamide, and two L-ascorbic acid molecules. Six D-glucose molecules complete one Unit of the structure. Both Fe-S and cytochrome clusters, as well as dehydrogenases, ensure the continuous operation of the Unit. The synchronized function of the three-stoke, three twin-cylinder engine results in continuous energy, ATP, and H^+^ production. Eukaryotic cells are equipped with the SET-AG and the SET-OP; thus, they can live in an anoxic and oxygenized environment. It is hoped that the SET concept developed here will help the better understanding the way of action regarding the cancer treatment with Vitamin C or glucose deprivation.

## 1. Introduction

### 1.1 Anaerobic glycolysis, aerobic glycolysis, and oxidative phosphorylation

The function of the energy-generating system of cells is well known. However, the efficiencies of anaerobic glycolysis, aerobic glycolysis, and oxidative phosphorylation are different. They produce 2, 4, and 36 adenosine triphosphates (ATP)/mol^−1^ glucose, respectively. We know less about the composition and development of the structure responsible for energy production.

The eukaryotic cell was formed through an ancient cell’s symbiosis using aerobic glycolysis and a modern cell with oxidative phosphorylation [1]. Thus, the cellular energy system may have gradually evolved. We assume that the energy systems of different nature cells are made up of the same building blocks. Here, we describe the putative phylogenetic origin of these structures. Eukaryotic cells can also function under anaerobic conditions, ensuring the body’s ability to regenerate. The aerobic-anaerobic shift is enabled by the hypoxia-inducible factor (HIF) [2, 3].

### 1.2 Mitochondria contain vitamin C

Korth et al. supposed that two L-vitamin C (ascorbic acid, L-AA) molecules are in the NADPH pocket, presumably near the adenine binding site in the inner membrane of the mitochondria [4]. Their conclusion is based on molecular mechanistic docking computations.

### 1.3 Properties of the atoms of the protonated adenine molecule

Turecek and Chen investigated the features of the atoms of the protonated adenine molecule in the gas phase, water clusters, and bulk aqueous solution. They calculated that proton in the adenine ion 1^+^ undergoes fast migrations among positions N1, C2, N3, N10, N7, C8, and N9, which results in an exchange of hydrogen atoms before the loss of a hydrogen atom forming adenine cation radical at a 415 kJ mol^−1^ dissociation threshold energy [5]. We suppose that the proton displacement acts as a signal to determine electrons’ path migrating in the adenine molecule. Thus, according to the results of Turecek and Chen, the adenine molecule has two entering points for electrons (N1 and N7), while electrons might leave the molecule through the N10 atom [5] (Illustration 1).

**Illustration 1.**
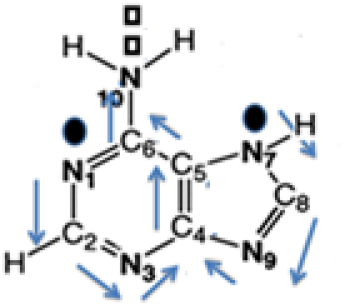
Two entering points (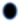 N1 and N7) and one leaving point (◻N10) of the adenine molecule’s electrons.

### 1.4 Fe-S clusters

Many Fe-S clusters are known in organometallic chemistry and as precursors to synthetic analogs of the biological clusters. The most abundant Fe-S clusters are of the rhombic [2Fe-2S], and cubic [4Fe-4S] types, but [3Fe–4S] and [4Fe–3S] clusters have also been described. Two probable structures for [Fe_2_S_2_(SCH_2_CH_2_OH)_4_]^e2-^ are presented in Illustration 2.

**Illustration 2.**
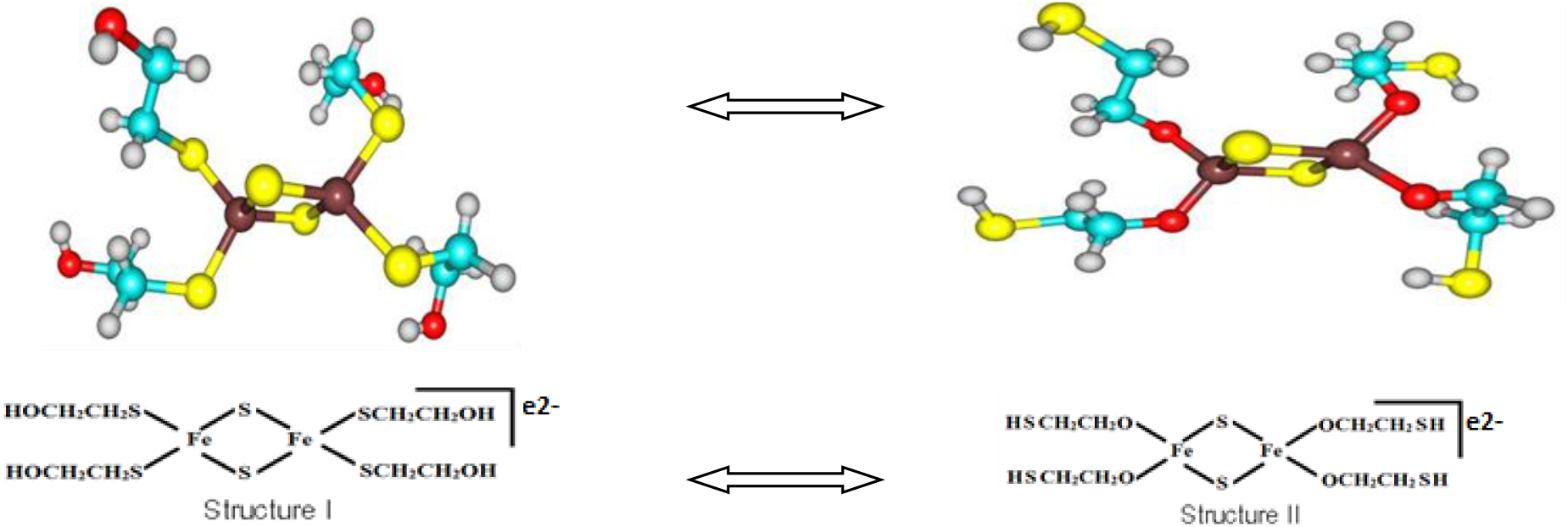
Two probable structures for [Fe_2_S_2_(SCH_2_CH_2_OH)_4_]^e2-^ and their density functional theory optimized structures. The structure I is when Fe coordinates SCH_2_CH_2_OH through S, while Structure II is when Fe coordinates SCH_2_CH_2_OH through the oxygen donor site.

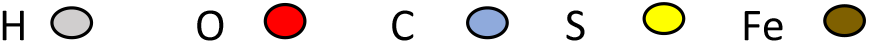

The properties of oxygen, sulfur, and iron allow the oxygen and sulfur atoms to swap places in the Fe-S cluster. The exchange is mediated by the electron flow facilitated by the nearby molecules.

### 1.5 ATP synthase forms rows of dimers in cristae membranes

The mitochondrial F1-FO ATP synthase is the most conspicuous protein complex in the cristae. ATP synthase is an ancient nanomachine. It uses the electrochemical proton gradient across the inner mitochondrial membrane to produce ATP by rotator catalysis [6, 7, 8].

### 1.6 Open questions

**We still do not know:**

1. How exactly Complex-I works, especially in which way electron transfer coupled to proton translocation happens?
2. Are there six or seven Fe-S clusters in Complex-I?
3. Where and exactly how the respiratory chain complexes are working?
4. From where does the ADP in Complex-V come?
5. How are the continuous electron transfer and ATP production managed?

## 2. Hypotheses

We assume that,

1. uric acid is not an end-product, but an intermediate molecule of the ATP - uric acid - ATP cycle, adenine molecules originate from the UA molecules,
2. the energy systems of different cells are made up of the same building blocks,
3. performed structures (SETs) are responsible for the energy transformation in the cells,
4. a tetra adenine octo phosphate ring (TAR) exists, in which the UA originated adenine molecules form a ring. The molecules are linked to each other through the N7-C2 and C8-N1 atoms of the TAR’s adenine molecules by H_2_PO_4_^e-^ molecules,
5. the four N10 atoms of the TAR’s adenine molecules bind one flavin, one nicotinamide, and two L-ascorbic acid molecules,
6. TAR + two L-ascorbic acids + nicotinamide + flavin + six D-glucose molecules create the ADP producing Unit (ADP-PU),
7. the ADP PU is the base element of all SETs,
8. one energy-producing twin motor of cells implements continuous production of energy, membrane potential, and ATP,
9. three (in SET-AG) or nine (in SET-OP) ADP-PUs results in the cells’ continuous electron transfer and energy supply.

### 2.1 ATP-uric acid-ATP cycle

The view that uric acid (UA) is the end-product of ATP metabolism is generally accepted. Gounaris et al. reported that the hypoxanthine molecule could develop as an effective capture of inorganic nitrogen species. These authors have also reported that hypoxanthine, the biochemical precursor of adenine and guanine, captured nitrite ions and was reductively transformed into adenine [9]. A similar reaction may occur by UA amination. We predict that UA is not an end-product but one element of an ATP-adenosine-inosine-hypoxanthine-xanthine-UA-aminated UA-adenosine-ADP-ATP cycle (Illustration 3).

**Illustration 3.**
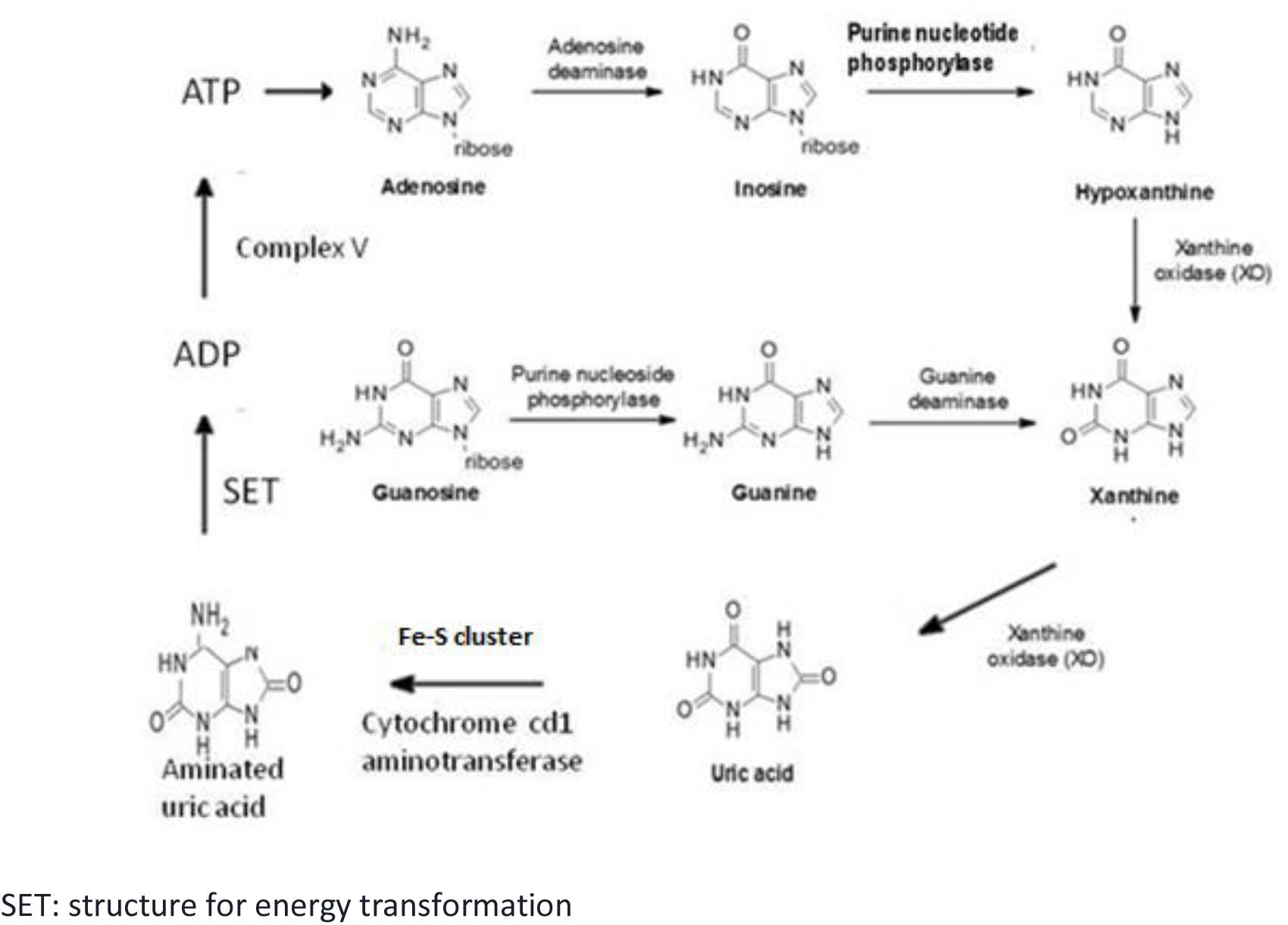
The hypothetical ATP-uric acid-ATP cycle. From the ATP, uric acid is created by the well-known metabolic way. Later, an aminated uric acid, adenosine, ADP, and ATP are generated from the uric acid.

#### Amination of uric acid

Fe-S clusters take up oxygen atoms from the C1 position of UA molecules. At the same time, UA will be nitrified with the help of the aminotransferase enzyme. For example, the 2Fe-2S cluster obtains four oxygen atoms from four UA molecules. Thus, they produce four aminated UA molecules and four H_2_O molecules.

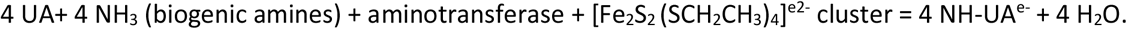

### 2.2 Hypothetical structure for energy transformation

Maintenance of life requires continuous electron flow. Cells provide this through permanent glycolysis resulting in continuous membrane potential, energy, ATP molecules, CO_2_, and H^+^. During evolution, the essential elements of the energy supply system were formed in the periplasm of the cell. These elements, like the molecules encoding the genetic trait, are fundamental structures of the living world.

We assume that, in every cell, SETs provide the cell with the proper energy supply and H^+^. Therefore, we think that the units of the SET consist of the same building blocks.

Our hypothesis is based on well-known data, such as the oxidative pentose pathway, the electron transport system in the mitochondria [10], the biochemical nature of H_2_PO_4_^e-^, and Fe-S clusters, the results of Turecek and Chen in connection with the nature of adenine [5], the oxidative experimental findings supported by stoichiometric computations, suggesting that there are two vitamin C molecules found in the NADPH pocket [4], and the known function of the mitochondrial F1-FO ATP synthase [6, 7, 8].

The JSME Editor, courtesy of Peter Ertl and Bruno Bienfait, was applied for calculating the presented three-dimensional (3D) structures of the hypothesized structures [11]. The figures primarily represent part of the structure under study. All pictures made by this program are signed with “JSmol.”

The base of our hypothesized ADP-PU consists of the FAD and NAD molecules and Complex V, completed with two adenine, two L-AA, and two D-glucose molecules.

According to our hypothesis, a unique SET is responsible for the continuous H^+^ and ATP production of living cells. We suppose that there are several versions of the SET. Two of them are described below. They are the SET of aerobic glycolysis (SET-AG) and the SET of oxidative phosphorylation (SET-OP).

#### Evolution of the energetic system in the living world

We suppose that the enzyme aminotransferase may create aminated UA and 4 PO_3_^e2-^ molecules with the help of Fe-S clusters and dehydrogenases. The question is whether nitrogen is coming from NO_2_^e-^, as suggested by Gounaris et al. [9], or from NH_3_, or biogenic amines.

### 2.3. Structures for energy transformation

#### 2.3.1. The nest of structure for energy transformation

We suppose that in the mitochondrial cristae of eukaryotic cells and the periplasm of all kinds of cells, sites (nests) formed by six Fe_2_-S_2_ clusters exist for energy transformation.

##### Mitochondrial nest

The nest’s position is identical to the Complexes I, II, III, IV, and V positions within the inner membrane [12]. Base molecules arrive at the nest, where successive production of ADP, ATP, and H^+^ molecules start. When the energy transformation is completed, ATP, CO_2_, and H^+^ leave the nest, and new base molecules will arrive. Thus, the nest is the determining frame of all SETs. We suppose that the three Fe-S clusters of Complex II are identical to the Fe-S clusters of Complex I.

##### Nest of bacterial ancestor cells

SET-AG is in the periplasm of the bacterial ancestor cells.

##### Eukaryotes

Eukaryotic cells have both SET-OP and SET-AG. However, SET-AG is in the periplasm, while SET-OP in the mitochondria.

#### 2.3.2. Adenine binding place

We suppose that the known NADPH pocket contains four adenine molecules. They are bound to each other by eight H_2_PO_4_ molecules through the N7-C2 and C8-N1 atoms of the adenine molecules. They form the TAR. Adenine is created from aminated UA; thus, the ring can be best described as a tetra-aminated UA ring at the start of energy transformation (Illustration 4).

**Illustration 4.**
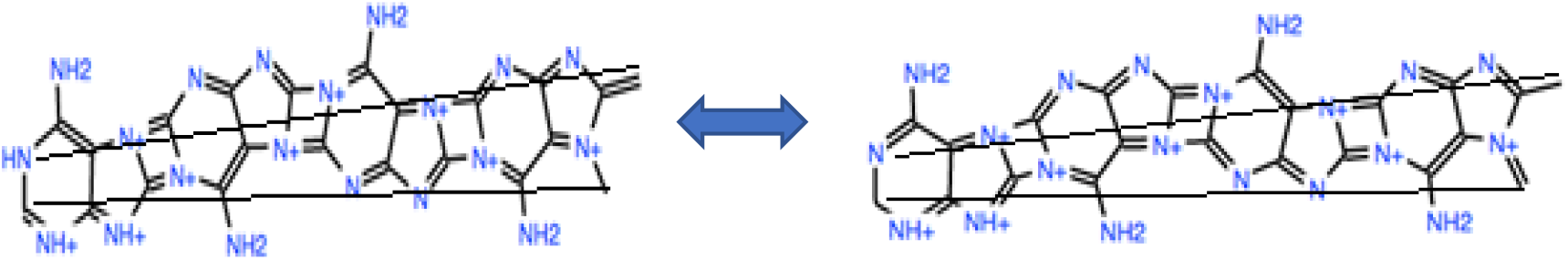
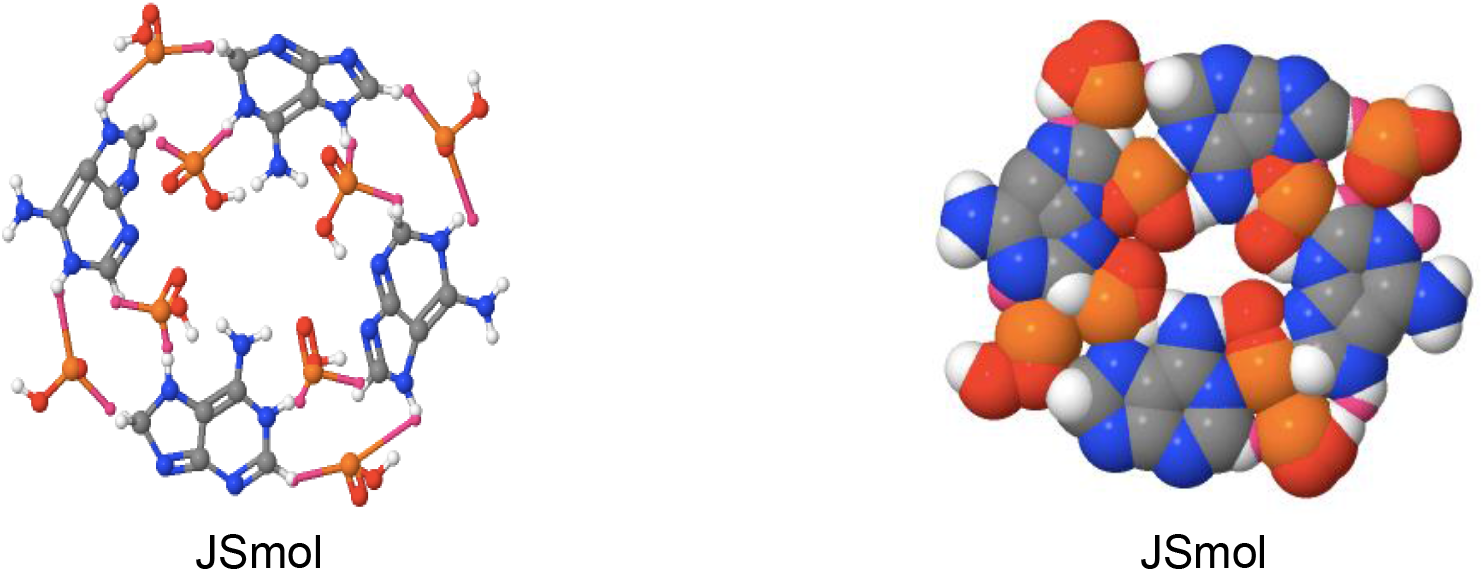
Tetra-adenine octo phosphate ring

The TAR can facilitate electron transfer. According to our hypothesis based on Turecek et al. [4], the N1 and N7 atoms of the adenine molecules are the entering points, while the N9 and N10 atoms might be the exit points for the electrons. The electron capture points are linked by H_2_PO_4_^e-^ molecules. In the TAR, electrons are arriving from the H_2_PO4^e-^ molecules (2^e-^).

#### 2.3.3. Molecules and units of the structures for energy transformation

##### 2.3.3.1. Determining molecules of the structures for energy transformation

The SET is made up of structural and support molecules. Fe-S clusters, UA-originated adenine, nicotinamide, flavin, L-Ascorbic-acid, D-glucose, H_2_PO_4_^e-^ and biogenic amines and cytochromes are the determinant molecules of all SETs. Fe-S cytochrome clusters, nicotinamide, and flavin are the constant frame (structure molecules), while UA, L-AA, D-glucose, H_2_PO_4_^e-^ and biogenic amines are continuously arriving at the SET, while ATP, CO_2_ H_2_O, and H^+^ are leaving it.

##### 2.3.3.2. Basic units of structures for energy transformation

###### The ADP Producing Unit, the Electron Flow Device

All SETs consist of the ADP-PUs. The ADP-PU contains six Fe-S clusters complemented with dehydrogenases, membrane transporters, and mitochondrial multienzyme complexes. In addition, four UA-originated adenine, one nicotinamide, one flavin, two L-AA, and six D-glucose molecules allow ADP-PU to work. The ADP-PU organizes the multienzyme complexes into functional units.

The ADP-PU consists of six [Fe_2_S_2_ (SCH_2_CH_3_)_4_]^e2-^ clusters. Four of the clusters are responsible for the oxidation of the 6^th^ carbon atoms of six D-glucose molecules and the two L-AA molecules. The 5^th^ [Fe_2_S_2_ (SCH_2_CH_3_)_4_]^e2-^ cluster oxidizes the 5^th^ carbon atoms of the two L-AA molecules, while the 6^th^ cluster, together with the aminotransferase enzyme and dehydrogenase, is responsible for the catalysis of the aminated UA molecules and the production of 4 PO_3_^e3-^.

All the SETs contain the ADP-PUs. The characteristics of supplementer molecules determine the nature and products of the structure.

###### SET of aerobic glycolysis

The SET of aerobic glycolysis consists of three ADP-PUs, each containing six [Fe_2_S_2_ (SCH_2_CH_3_)_4_]^e2-^ clusters and 3 Complex V. Accordingly, it can produce 3 × 4 ATP molecules.

###### SET of oxidative phosphorylation

The SET-OP consists of nine (3×3) ADP-PUs. It also contains one pyruvate dehydrogenase complex (PDC) and three high molecular weight cytochromes (Hmc). Thus, SET-OP produces 36 ATP molecules.

Other quantities of Fe-S and cytochrome clusters can result in additional SETs.

The SET-AG is situated in the cells’ plasma membrane, while the SET-OP is in the mitochondrial cristae.

#### 2.3.4. The reaction of the Fe-S cluster with aminated uric acid and H_2_PO_4_^e-^

In each of the four connecting points of the TAR, two aminated UA molecules and two H_2_PO_4_^e-^ molecules react with one Fe-S cluster. We believe that the ADP-PUs use aminated UA molecules. Aminated UAs arrive at sites defined by four Fe-S clusters (Illustration 5). Illustration 5A shows the nest of the ADP-PU before the start, and Illustration 5B shows when oxygen atoms (O^e2-^, circled) are already prepared for the oxidation of carbon atoms. The glucose molecules are not illustrated.

**Illustration 5.**
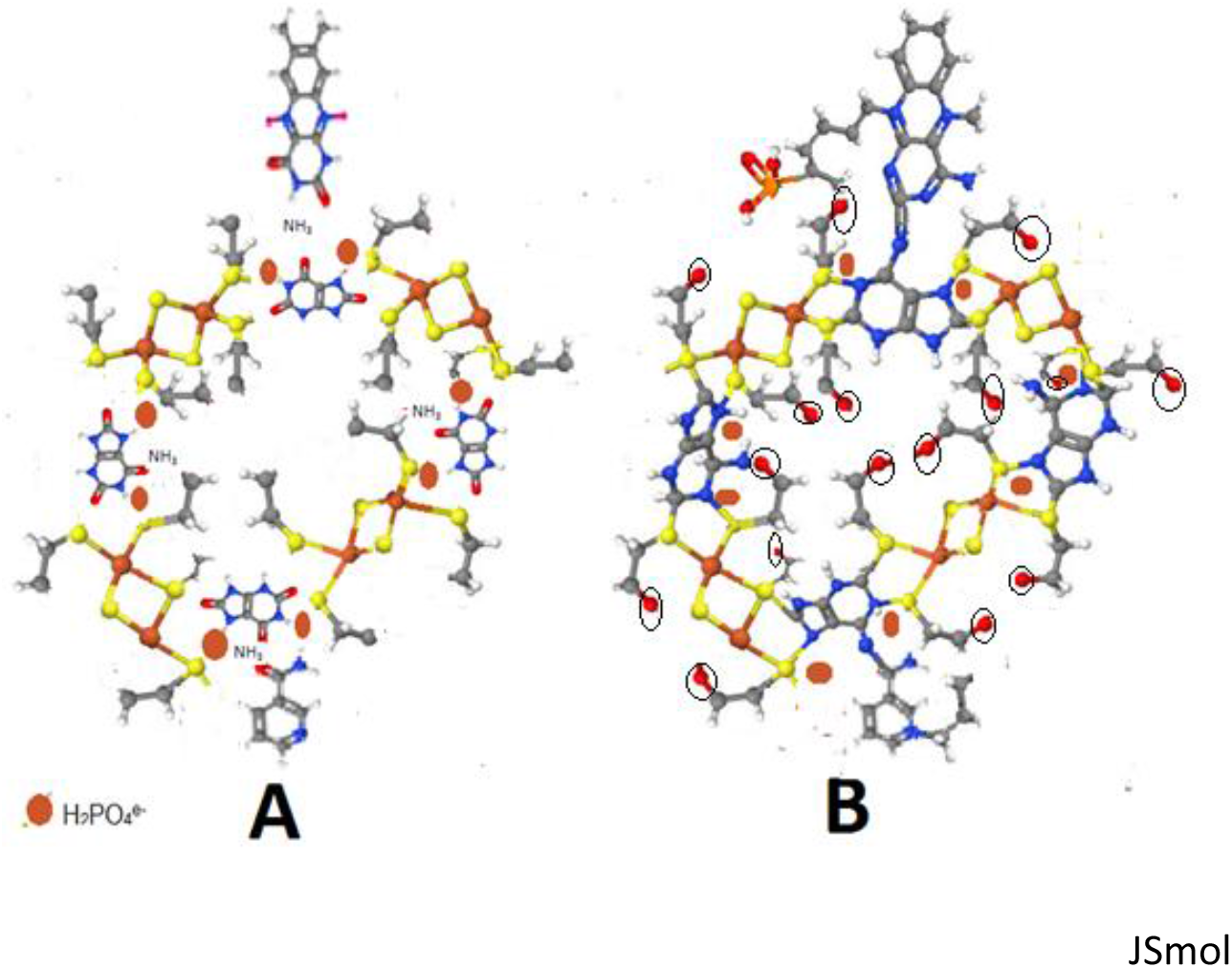
A: Four [Fe_2_S_2_ (SCH_2_CH_3_)_4_]^e2-^ clusters, flavin, and nicotinamide molecules are waiting for the supplementer molecules (eight H_2_PO_4_, four NH_3_ [biogenic amines], and four uric acids). The D-glucose and L-ascorbic acid molecules are not presented. B: Oxygen atoms (O^e2-^, circled) of four aminated uric acid molecules and eight H_2_PO_4_/PO_3_ molecules are ready to oxidate carbon atoms. One D-glucose molecule is presented.

In each of the four Fe-S clusters, the sulfur atoms are replaced by oxygen-derived from H_2_PO_4_^e-^ (eight oxygen atoms) and aminated UA molecules (eight oxygen atoms).

One of the four connecting points is presented in Illustration 6, with arrows indicating the supposed electron flow. In each connecting point of the TAR, two electrons are transferred. I think that when adenine molecules are organized into the TAR, the electrons pass the molecule not only in the direction of the N10 atom, as presented in Illustration 1, but also in the direction of the N9 atom (Illustration 6). The 3D picture is available at https://set.suicidevolution.com/v2/

**Illustration 6.**
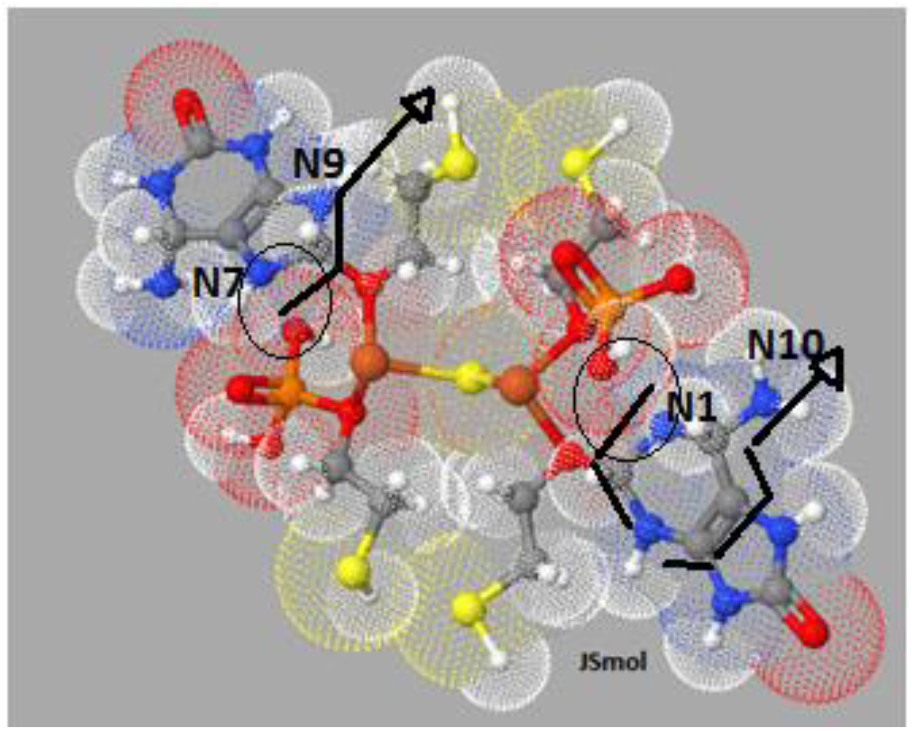
Two aminated uric acid molecules and two H_2_PO_4_ molecules arrive in one [Fe_2_S_2_ (SCH_2_CH_3_)_4_]^e2-^ cluster. The aminated uric acid molecules and H_2_PO_4_^e-^ molecules are bound in the Fe-S cluster. H_2_PO_4_^e-^ is not yet bound to the aminated UA (circle). Arrows indicate the supposed electron flow.

As a result, the tetra UA/adenine ring is formed. Its interfaces have electrons entering and leaving points. The H_2_PO_4_^e-^ molecule, bound by oxygen through the Fe-S cluster, is located near an electron entering point (circle) that allows electrons to enter the adenine molecule (Illustration 6). At each entry point, two electrons coming from two hydrogen atoms will leave behind two protons. Finally, the four UA-originated adenine molecules and the eight H_2_PO_4_ molecules form one TAR.

#### 2.3.5. Process of uric acid-adenine conversion during energy transformation

Four aminated UA molecules arrive in four of the six Fe-S clusters of the ADP-PU, where the UA molecules’ oxygen atoms are bound by four of the iron-sulfur complexes. The Fe-S clusters also bind eight H_2_PO_4_^e-^ molecules to one of their oxygen atoms.

In the next step, eight H_2_PO_4_^e-^ molecules are bound to the newly formed four adenine molecules, linking their N7-C2 and C8-N1 atoms. Subsequently, 8 × 2 electrons arrive at the TAR via the adenine molecules’ electron-accepting points (N1 and N7), leaving them at the N9 and N10 atoms. The Fe-S cluster’s connection with UA forms the possibility of electron transfer in the direction of the nitrogen atoms (N9 and N10) of the adenine molecules (Illustration 6).

#### 2.3.6. Nicotinamide, flavin, and vitamin C molecules connected to tetra adenine ring Amide-carbonyl hydrogen bond

The amide-carbonyl hydrogen bond determines the proper function of the SETs, as all ADP-PUs consist of four amide-carbonyl connections (flavin-adenine, nicotinamide-adenine, and two L-AA-adenine). Accordingly, dehydrogenases are important protein structures catalyzing the reactions at the connecting points of the ADP-PUs.

The N10 atoms of the aminated UA molecules connect to additional molecules. The supposed binding to flavin, nicotinamide, and L-AA molecules is presented in Illustration 7. The tetra adenine ring binds nicotinamide, flavin, and two L-ascorbic acid molecules with the help of dehydrogenases. The double bond between the N10 atoms of adenine and these molecules allows the electron flow. It provides rapid connection and separation of these molecules in the TAR (Illustration 7).

**Illustration 7.**
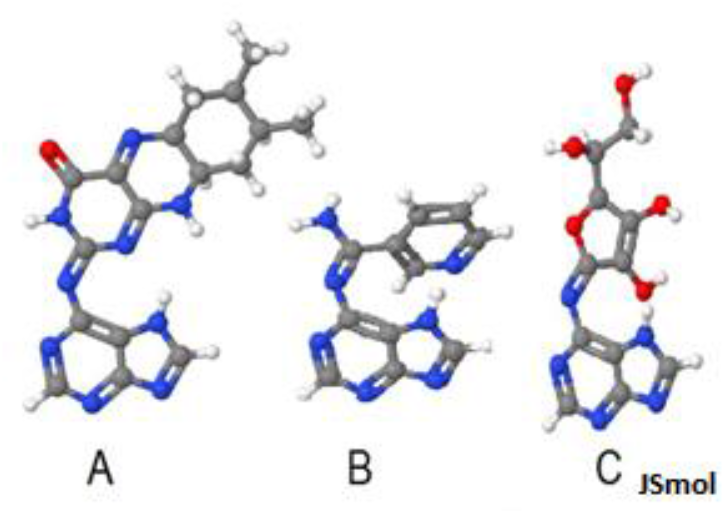
Presumed interfaces of adenine-flavin (A), adenine-nicotinamide (B), and adenine- L-ascorbic acid (C) molecules.

The 5th and six^th^ [Fe_2_S_2_ (SCH_2_CH_3_)_4_]^e2-^ clusters are responsible for the oxidation of the 5th carbon atom of the two L-AA molecules, for the amination of the 4 UAs, and the production of 4 PO_3_^e3-^.

#### 2.3.7. The synthesis of adenosine

Before the electron flow starts, the Unit is completed by six D-glucose molecules (Illustration 8). Four D-gluconate molecules (1–4) are bound to the adenine molecules via a β-N9-glycosidic bond (Illustration 8A), one D-glucose (5) to the nicotinamide, and one (6) to the flavin molecule (Illustration 8B).

**Illustration 8.**
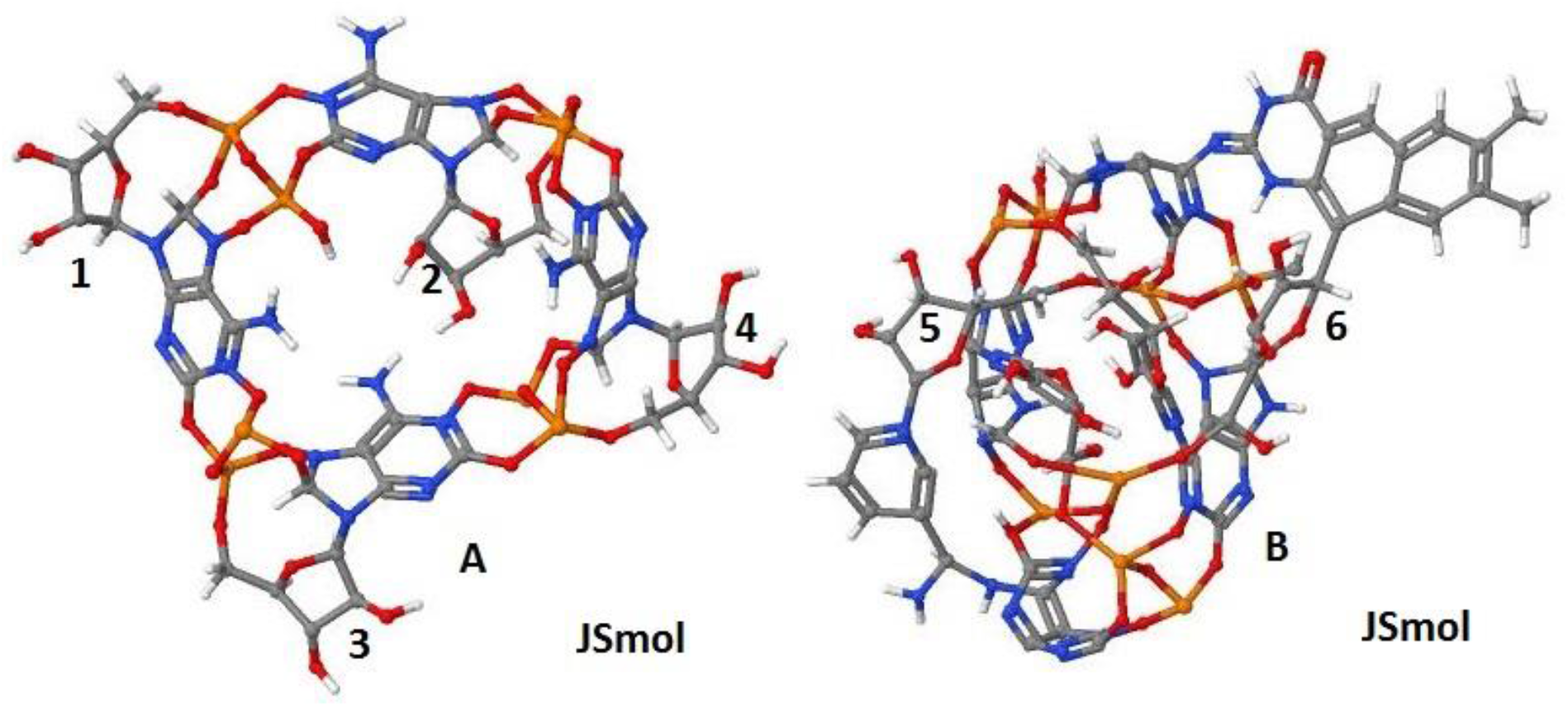
Six D-glucose molecules in one Unit of the SET in the tetra adenine octo phosphate ring. Pictures A and B illustrate the same Unit of the SET. Picture A presents four 5-phospho-D-gluconate molecules (1–4), and picture B presents two other 5-phospho-D-gluconate molecules (5, 6), along with one flavin, one nicotinamide, and two ascorbic acid molecules. The 6^th^ carbon atoms of the glucose and L-AA molecules are oxidized (not presented). Accordingly, the picture shows ribose molecules created from the D-Glucose molecules. https://set.suicidevolution.com/v2/

#### 2.3.8. Rebirth of vitamin C

Nicotinamide, flavin, and vitamin C molecules in the ADP-PUs

The location of the flavin mononucleotide molecule within one ADP-PU is shown in Illustration 9A. One adenine (a) will be bound to flavin, which will also bind one 5-phospho-D-gluconate molecule. An additional 5-phospho-D-gluconate molecule will bridge this structure to another adenine (c) via a β-N9-glycosidic bond (not presented).

**Illustration 9.**
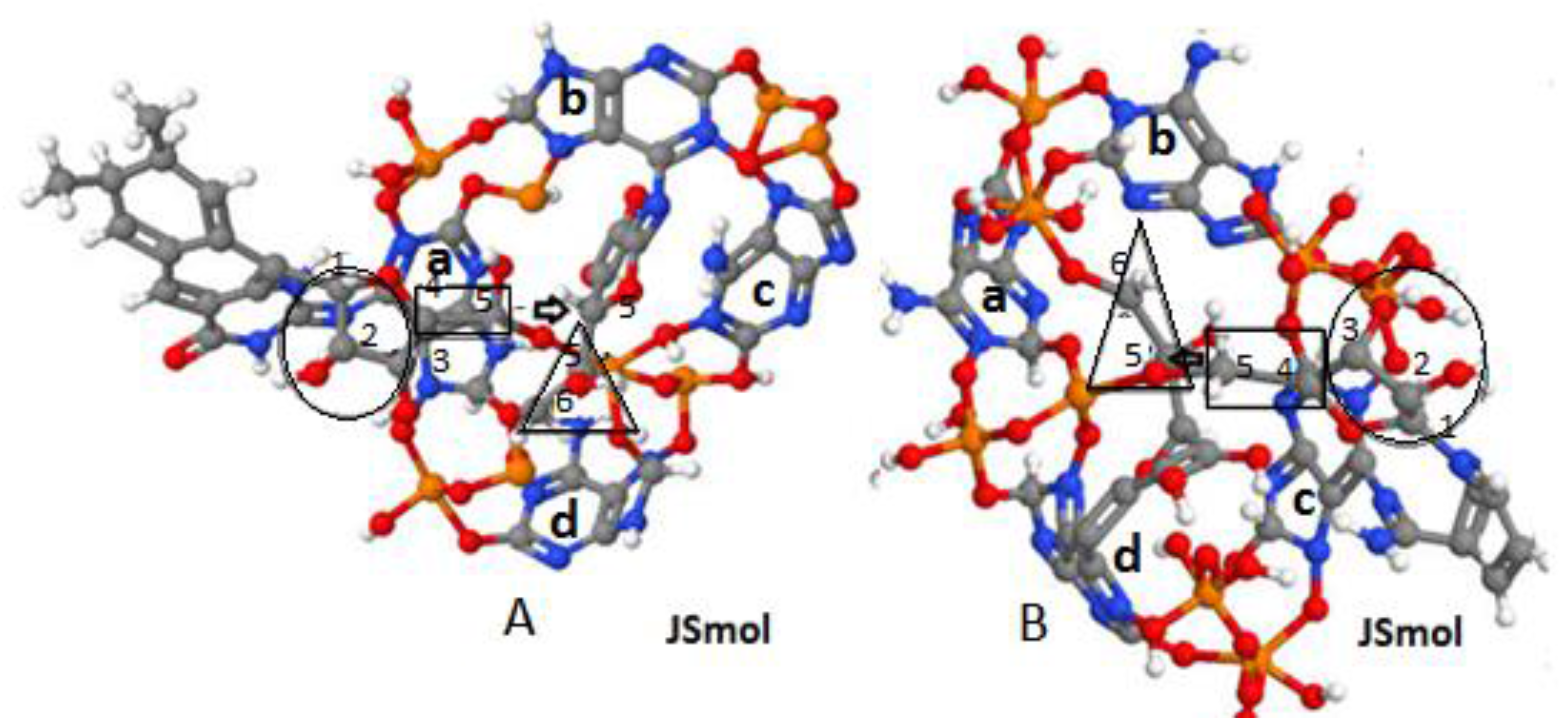
Two pictures of the same Unit (A and B). A: Flavin riboside molecule and one L-vitamin C molecule in the tetra adenine octo phosphate ring. B: Nicotinamide riboside molecule and one L-vitamin C molecule in the tetra adenine octo phosphate ring. The 5^th^ and 6^th^ carbon atoms of the two L-AA molecules will be oxidized (triangle). Two × 2 carbons (quadrilateral) will be connected to the L-AA molecules’ remaining two lactone rings (arrows). Two pyruvate molecules are situated in the two circles.

The location of the nicotinamide riboside molecule within one ADP-PU is shown in Illustration 9B. One adenine (c) will be bound to nicotinamide, which will also bind one 5-phospho-D-gluconate molecule. An additional 5-phospho-D-gluconate molecule will bridge this structure to another adenine (a) via a β-N9-glycosidic bond (not presented).

The 1^st^, 2^nd^, and 3^rd^ carbon atoms of the 5-phospho-D-gluconate molecules bound to flavins and nicotinamide will create two pyruvate/lactate molecules (circle) (Illustration 9).

The 5^th^ and 6^th^ carbon atoms of the two L-AA molecules will be oxidized (triangle). Then, two × 2 carbon atoms (quadrilateral) will be connected to the remaining two lactone rings of the L-AA molecules (arrows), forming two renewed L-AA molecules (Illustration 9). The 3D versions of the pictures are available at https://set.suicidevolution.com/v2/.

In an oxygenated environment, pyruvate molecules of NAD and FAD will be oxidized by the PDC. Accordingly, in an anoxic environment, 9 × 10, and in an oxygen-containing milieu, 9 × 16 carbon atoms, respectively, will be oxidized in the nine ADP-PUs of SET-OP.

### 2.4. The products of the ADP-PU

In all the SET units, the disconnected carbon atoms will be oxidized by eight 2O^e2-^ delivered by the four Fe-S clusters. At the same time, an additional Fe-S cluster prepares two 2O^e2-^ responsible for the oxidation of the 5^th^ carbon atoms of the two L-AA molecules.

The products are four ADP, two ribose (which will be converted to two pyruvate and two citric acid molecules), two lactone rings, 10 H_2_O, 10 CO_2_, and 20 H^+^. In an anoxic environment, the pyruvate will be conversed to lactate, while in the presence of O_2_, it will be oxidized to three CO_2_. In addition, the lactone rings will bind the citric acid molecules, resulting in two renewed L-AA molecules. Thus, the end-product of the reaction is four ADP, two pyruvates, 10 CO_2_, two renewed L-AA molecules, 4 PO_3_^e3-^ and 20 H^+^.

### 2.5. Continuous electron and ATP production

#### 2. 5. 1. The three steps of ATP production

In all the SETs, ADP-PU, TAR, Hmc, and cytochrome C work in close cooperation. Besides, the connections of the three nicotinamide and three flavin molecules result in the cooperation between the three ADP-PUs (Illustration 10). In this way, one of the units is permanently active. Moreover, the coordinated operation of the three subunits ensures continuous electron, ADP, and ATP production. Thus, the SET-AG has three ADP-PUs, while the SET-OP has nine (3 × 3) ADP-PUs.

**Illustration 10.**
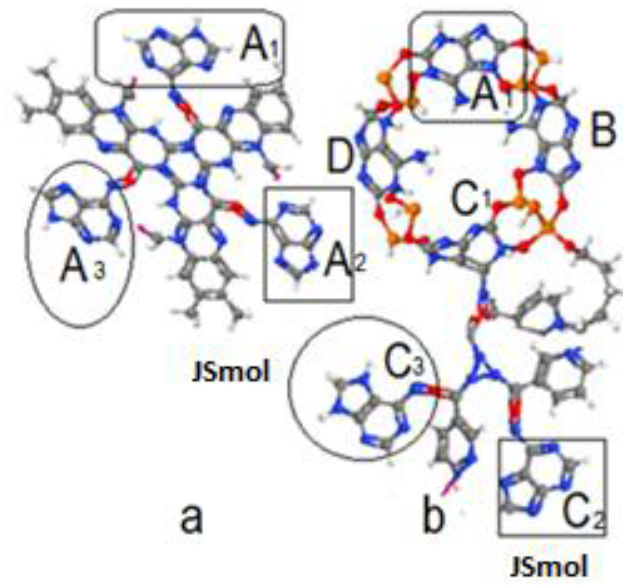
a: three flavins + three adenines. b: three nicotinamide molecules, two adenine molecules, and one tetra adenine octo phosphate ring. A_1_ of the picture a is identical to the A_1_ of picture b. A_2_ interacts with C_2_, while A_3_ relates to C_3_ by two adenine molecules, creating two other tetra adenine octo phosphate rings.

Three flavin molecules bind three adenine molecules (A) on the surface of the inner membrane facing the cytoplasm or matrix (Illustration 10a), and 1.5 nm away, three nicotinamide molecules bind three adenine molecules (C) in the inner membrane so that the adenine molecules (C) are superimposed on the flavin-bound ones (A). Thus adenine molecules A and C form a TAR using two additional adenines (B and D) (Illustration 10b).

We suppose that the three ADP-PUs’ synchronized operation allows continuous electron and ATP production, as one Unit is permanently active, and the functions of Complex V and the ADP-PUs are also synchronized (Illustration 11). When the ADP-PU is in an open state (releasing ADP), Complex V is loose (accepting the ADP molecules).

**Illustration 11.**
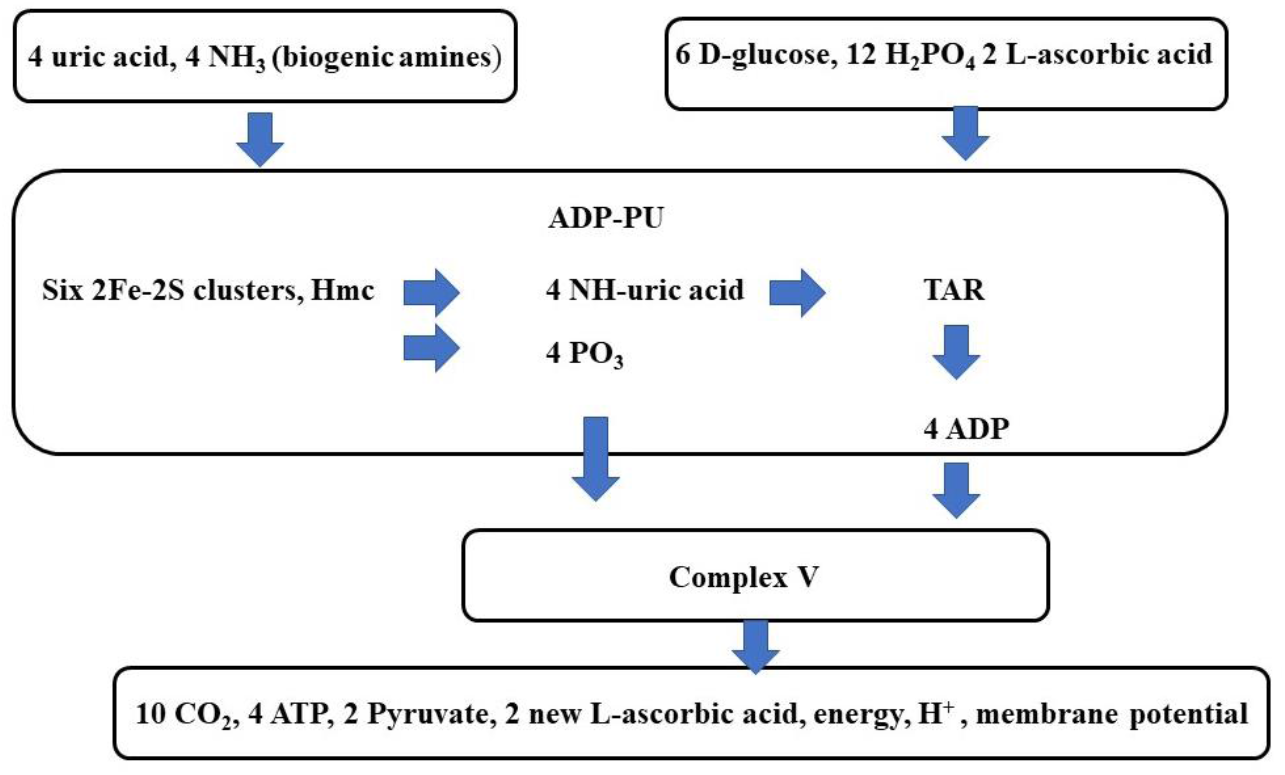
Cooperation between Fe-S, high molecular weight cytochrome (Hmc) clusters, and Complex V in the Structures of Energy Transformation.

#### 2.5.2 The process of energy transformation

In the first step, Hmc or Fe-S clusters produce aminated UA and PO_3_^e3-^ molecules. The aminated UA molecules will form TARs. The second step of energy transformation is ADP production by the ADP-PU. In the third step, ATP synthase (Complex V) will finish the reaction, resulting in the formation of ATP from ADP + PO_3_^e3-^. In the SET-OP, the PDC and Hmc clusters catalyze the conversion of pyruvate molecules to CO_2_. H^+^ is liberated during the reaction catalyzed by the ADP-PU. Protons created by Hmc clusters in the cristae space flow back into the matrix through the ATP synthase rotor, driving ATP production (Illustrations 11).

### 2.6. The supposed complete structures for energy transformation

The SET may consist of the following units: ADP-PU, and functional elements, such as Complex V, PDC, and Hmc clusters.

## 3. The energy-producing twin motor of cells

All cells are powered by one synchronous, three-stroke twin-engine system. The system operates by the well-known clusters, complexes, and structural proteins. Complex V and ADP-PU make up the twin engine. The coordinated operation of three ADP-PUs and Complex V ensures that, in the SET-AG, one of the three units (or three of the nine ADP-PUs of the SET-OP) is always active, resulting in the continuous membrane potential and ATP production.

### Operating phases of the three-stroke energy converter nano engine

*The first (loose) phase*:

ADP-PU: Essential (parent) molecules enter ADP-PU.
Complex V: ADP and PO_3_^e3-^ enter Complex V.

*The second phase (active state)*:

ADP-PU: Enzymes and 2F-2S clusters transform the parent molecules to form ADP and PO_3_^e-^ molecules.
Complex V: ATP is formed from ADP and PO_3_^e3-^.

*The third phase (open state):*

ADP-PU: ADP and PO_3_^e-^ leave the ADP-PU.
Complex V: ATP leaves Complex V.

When the ADP-PU is in the open state, releasing ADP and PO_3_^e-^, Complex V is in a loose state, ready to uptake ADP and PO_3_^e-^ (illustration 12).

**Illustration 12.**
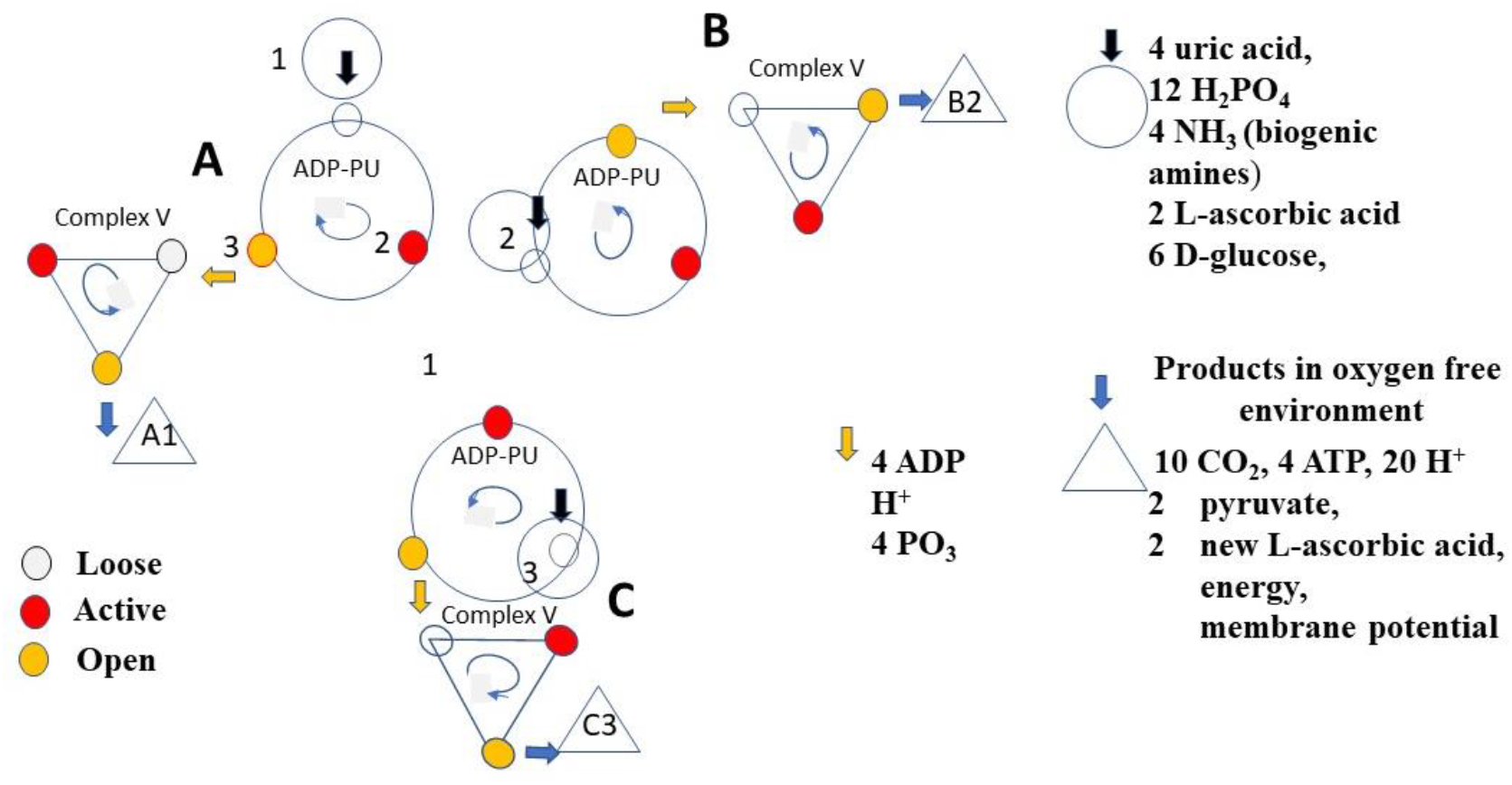
The synchronized function of the three-stroke, three twin-cylinder (A, B, C) engine of the ADP-Producing Unit.

## 4. The size of the structures for energy transformation

We calculated the sizes of the SETs using the JSME Editor courtesy of Peter Ertl and Bruno Bienfait [11]. Based on these data, we estimate the SET-AG as a sphere with a 1.5 nm diameter. The SET-OP consists of 9 (3×3) ADP-pus. Thus, most probably, it is one 2.0–3.0 nm spherical structure (Illustration 13).

**Illustration 13.**
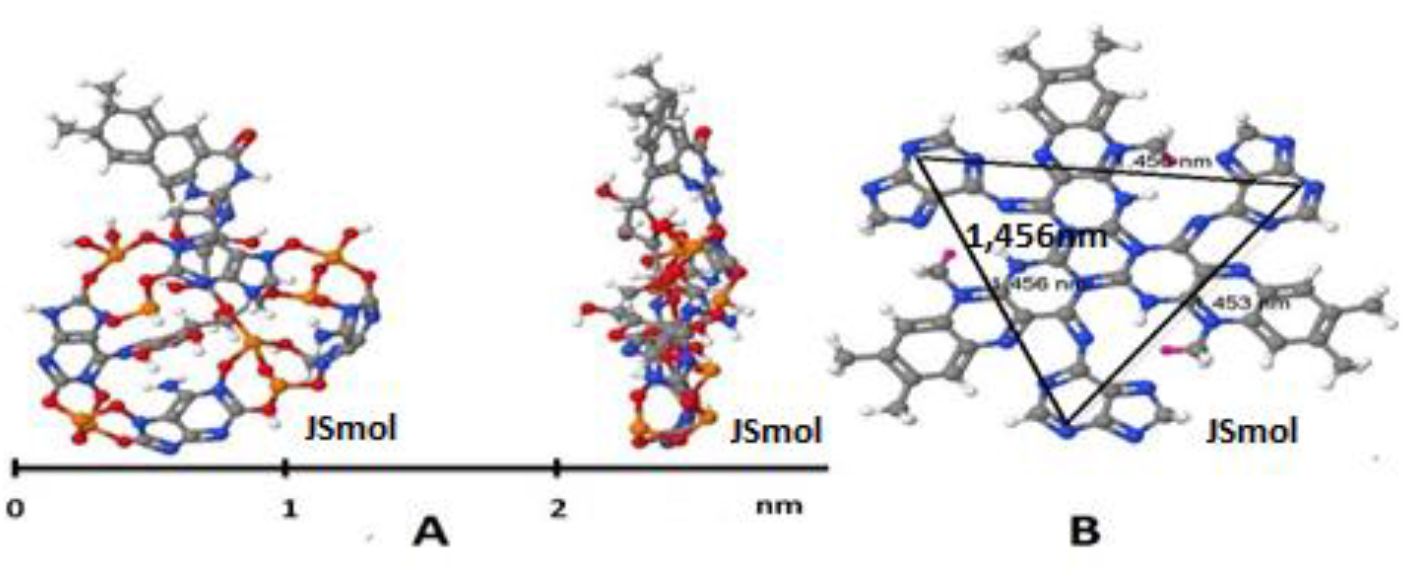
The size of the structure for energy transformation’s units. One tetra adenine octo phosphate ring + one vitamin C + one flavin + one glucose molecules (A) and the three units’ distance (B).

## 5. The eukaryotic cells*’* dual energetic supply

### 5.1 Symbiosis of one ancient cell with a new cell resulted in eukaryotes

Life was created in an ancient world without O_2_. Accordingly, the defenses of the first living cells against free radicals were low. Thus, the emergence of O_2_ produced by cyanobacteria resulted in an environmental disaster, and most of the living cells died. Simultaneously, cells were created that could produce significantly more energy using O_2_ than ancient cells. In addition, the new cells were armed with an effective defense system against free radicals [13].

Lynn Margulis (1938-2011) thought that the eukaryote cells’ ancestors avoided being destroyed by oxygen by entering a symbiotic relationship with aerobic bacteria (illustration 14) [14]. Accordingly, mitochondria are responsible for breathing and the effective defense against free radicals in the eukaryotic cells. Mitochondria provide the cell with a significantly better energy supply and free radical protection. Eukaryotic can function in both anoxic and oxygenized environments [14].

**Illustration 14.**
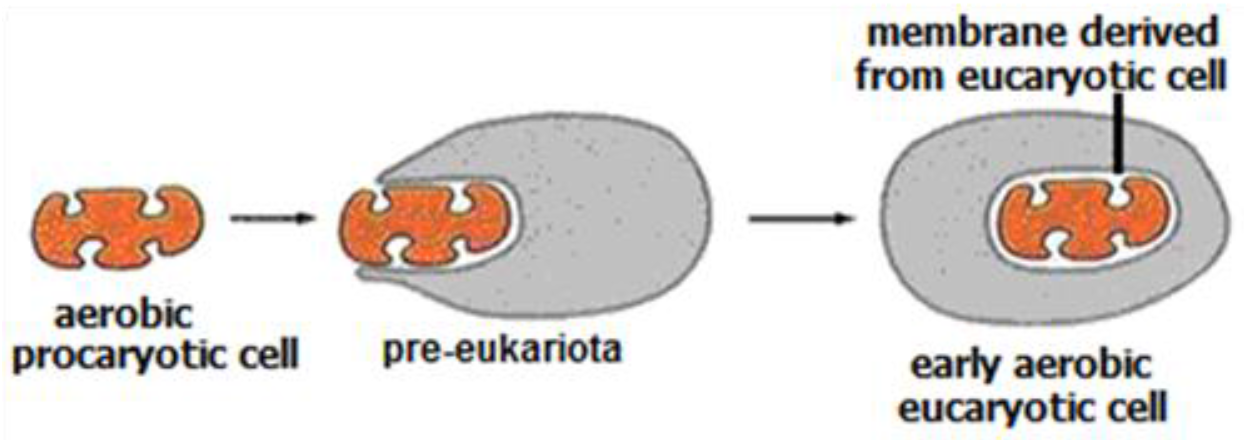
The evolution of mitochondria according to the theory of endosymbiosis.

### 5.2. The HIF system

The HIF system is the detector and conductor of the oxygenated and oxygen-free environment. Therefore, it can facilitate the cell back to ancient times.

The HIF system ensures adaptation to an environment without O_2_ ([2, 3, 15, 16]. In an anoxic environment, cells use aerobic glycolysis offered by the SET-AG. In the existence of O_2,_ the SET-OP presents oxidative phosphorylation (Illustration 15).

**Illustration 15.**
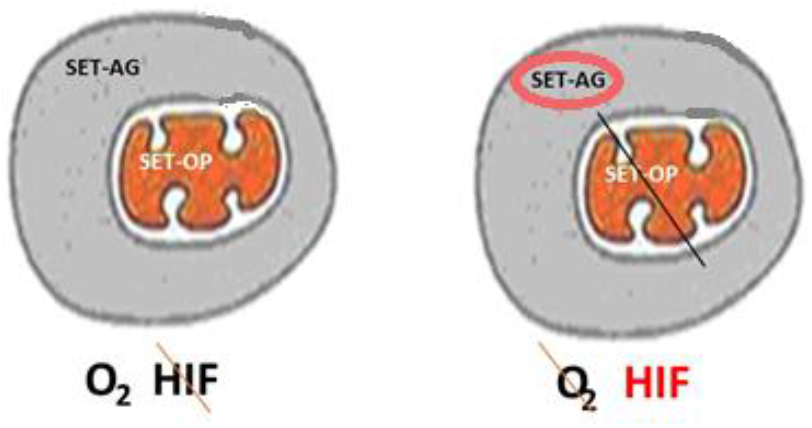
The HIF system is the detector and conductor of the oxygenated and oxygen-free environment. **Abbreviations:** SET-AG: Structure for Energy Transformation of Aerobic Glycolysis; SET-OP: Structure for Energy Transformation of oxidative phosphorylation; HIF: Hypoxia-Inducible Factor.

### 5.3. Warburg theory of cancer

Warburg postulated that the driver of tumorigenesis is insufficient cellular respiration. He hypothesized that cancer’s malignant growth is caused by the fact that tumor cells mainly generate energy (e.g., adenosine triphosphate / ATP) by the breakdown of glucose (a process called aerobic glycolysis) in contrast with healthy cells that mainly generate energy from pyruvate’s oxidative breakdown (oxidative phosphorylation) [17 – 19].

### 5.4. Advantages of the symbiosis

1. The energy efficiency of the eukaryotic cell is significantly better than that of the ancient cell. For example, mitochondria can produce 36 ATP/mol glucose through oxidative phosphorylation, while the ancient cell has an efficiency of 3 ATP/mol glucose.
2. Effective mitochondrial defense system against free radicals.
3. Function in both anoxic and oxygenized environments.

In the body formed by eukaryotes, cells can adapt to the O_2_ deficiency. Due to the lack of O_2_ caused by injury or any cause, the hydrolysis of HIF-1α is canceled. As a result, the HIF system influences about 200 genes, converting the cell to ancient energetics.

The most significant changes are:

- Ensuring the cell’s increased glucose demand due to the ancient energy system’s lower efficiency.
- Reduction of the apoptosis threshold.
- Induction of neovascularization, adhesion molecules, and stem cells.

As a result, the circulation will be restored with the help of newly formed blood vessels. In addition, increasing tissue O_2_ will hydrolyze the HIF-1α; thus, the cells will return to the mitochondrial oxidative phosphorylation.

### 5.5. Disadvantages of the symbiosis

In the case of malignant tumor development, the cancer cells will multiply without control. Proliferating cells gradually move further away from the vessel supply; thus, an anoxic area will be formed since there is no vasculature in the newly developing tumor. With the HIF system’s help, tumor cells switch to the ancient energy system, creating the cancer cell characterized by Warburg’s aerobic glycolysis. The HIF system will ensure the cell’s increased glucose demand, reduce the apoptosis threshold, induce neovascularization and adhesion molecules allowing the evolution process of cancer [20]. As a result of this process, a tumor is created by cells using original oxidative phosphorylation and cells with Warburg’s aerobic glycolysis. Tumor cells using Warburg’s aerobic glycolysis are more sensitive to free radicals than the normal cells, and the tumor cells using oxidative phosphorylation as the effective defense against free radicals depends on the active mitochondrial function and the presence of O_2_.

## 6. Conclusions

The model of the SET helps to answer the questions described in the introduction. Ad 1: Complex I and the electron transfer chains work in a 3D structure of the EFD, responsible for the proton-electron translocation. Ad 2: the number of Fe-S clusters is six in the ADP-PU. 3: the ADP-PU and Complex V work is synchronized. Ad 4: the ADP-PUs produce ADP in all units of the SET. Ad 5: the continuous ATP production and membrane potential provided by continuous H^+^ production is realized by the synchronized operation of three or nine (3X3) ADP-PUs.

### 7. Possibilities for proving the hypothesis

The hypothetical SET model described above is a simplified idea. However, the SET is undoubtedly an extremely complex system with different modalities in different organisms.

We believe that experimental testing of the SET model is justified and warranted. If the model is correct, the following significant results will serve to confirm it:

1. Metabolism of ^13^C labeled D-glucose will yield ^13^C labeled L-AA in human or guinea pig cells. The change can be detected by nuclear magnetic resonance spectroscopy. Our unpublished preliminary data support the hypothesis.
2. ^13^C labeled citric acid might yield ^13^C labeled L-AA.
3. Delivery of ^13^C radiolabeled L-AA should result in *in vitro* and animal experiments in the appearance of labeled CO_2_ if the 5^th^ and or 6^th^ carbon atoms were labeled. In contrast, the CO_2_ labeling is omitted if the C1, C2, C3, or C4 atoms of L-AA were labeled.
4. Radiolabeled UA molecules will result in labeled adenine molecules.

**Table I.**
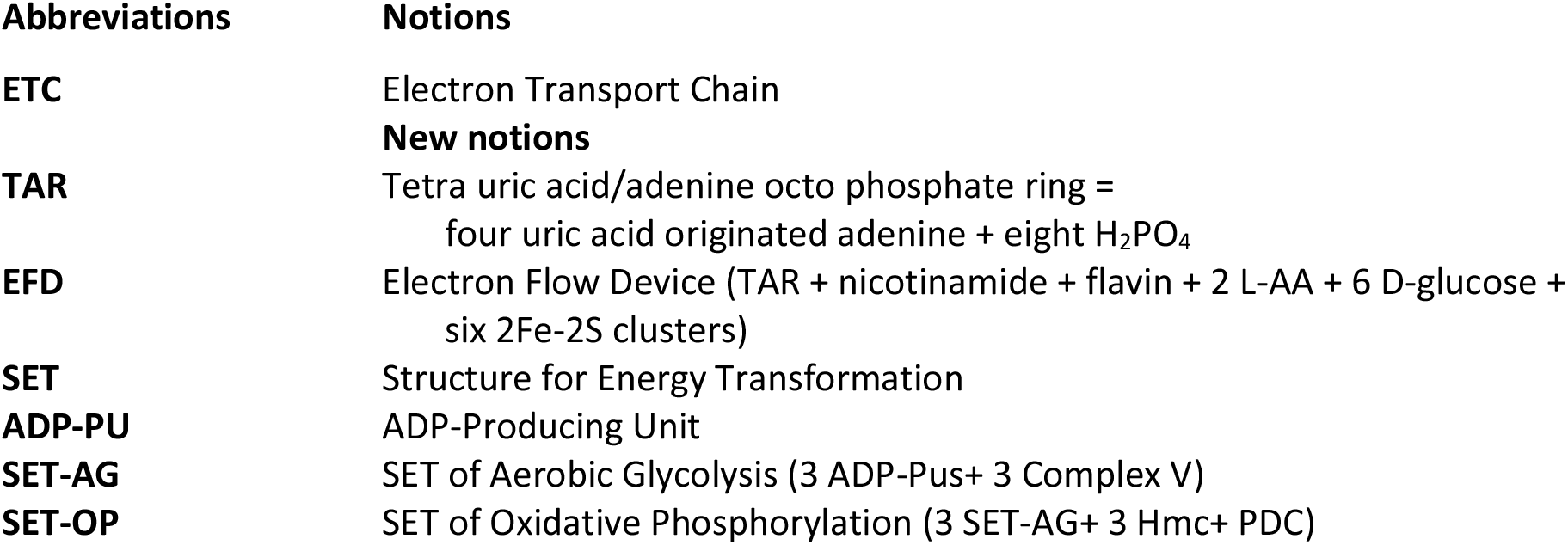
Notions and Abbreviations

## References

1. Margulis L Origin of eukaryotic cells New Haven: Yale University Press, 1970.

2. Rezvani HR, Ali N, Nissen LJ, Harfouche G, de Verneuil H, Taïeb A, et al. HIF-1α in epidermis: Oxygen sensing, cutaneous angiogenesis, cancer, and non-cancer disorders J Invest Dermatol, 2011;131:1793–1805.

3. Nauta T, van Hinsbergh VW, Koolwijk P Hypoxic signaling during tissue repair and regenerative medicine Int J Mol Sci, 2014;15:19791–19815.

4. Korth H, Meier A, Auferkamp O, Sicking W, de Groot H, Sustmann R, et al. Ascorbic acid reduction of compound I of mammalian catalases proceeds via specific binding to the NADPH binding pocket Biochemistry, 2012;51:4693–4703.

5. Turecek F, Chen X Protonated adenine: Tautomers, solvated clusters, and dissociation mechanisms J Am Soc Mass Spectrom, 2005;16:1713–1726.

6. Kühlbrandt W Structure and function of mitochondrial membrane protein complexes BMC Biol, 2015;13:10.1186/s12915-015-0201-x.

7. Hahn A, Vonck J, Mills DJ, Meier T & Kühlbrandt W (2018) Structure, mechanism, and regulation of the chloroplast ATP synthase. 2018;360:6389,4318; DOI: 10.1126/science.aat4318

8. Morales-Rios E, Montgomery MG, Leslie AGW, Walker JE Structure of ATP synthase from Paracoccus denitrificans determined by X-ray crystallography at 4.0 Å resolution PNAS, 2015;112(27, 43): 13231–13236 https://www.pnas.org/content/112/43/13231

9. Gounaris Y, Litinas C, Evgenidou E, Petrotos C A hypothesis on the possible contribution of free hypoxanthine and adenine bases in prebiotic amino acid synthesis Hypothesis, 2015;13: 10.5779/hypothesis.v13i1.393.

10. Michal G, editor Biochemie-Atlas Heidelberg: Spektrum Akademischer Verlag, 1999.

11. Ertl P, Bienfait B JSME Editor, In: http://biomodel.uah.es/en/DIY/JSME/draw.en.htm

12. Lee H, Yoon Y Mitochondrial membrane dynamics-functional positioning of OPA1 Antioxidants, 2018;7:10.3390/antiox7120186.

13. Cooper G.M. (2000). The Cell: A Molecular Approach. 2nd edition. 2000, Bookshelf Washington, DC: ASM Press; ID: NBK9841, Sunderland, Mass.: Sinauer Associates.

14. Lynn M. (1970). Origin of Eukaryotic Cells: Yale University Press. ISBN-10: 0300013531, ISBN-13: 978-0300013535 https://doi.org/10.1002/jobm.19730130220.

15. Huang L., Gu J., Schau M., Bunn H. (1998): Regulation of hypoxia-C 1alpha is mediated by an O_2_-dependent degradation domain via the ubiquitin-proteasome pathway. Proc Natl Acad Sci USA; 95(14):7987–7992. https://doi.org/10.1073/pnas.95.14.7987.

16. Knowles H., Raval R.R., Harris A.L., Ratcliffe P.J. Effect of ascorbate on the activity of hypoxia-inducible factors in cancer cells. Cancer Res, 2003;63:1764–1768. PMID: 12702559.

17. Warburg O. On the origin of cancer cells. Science, 1956;132:309–314 DOI: 10.1126/science.123.3191.309.

18. Alfarouk K., Verduzco D., Rauch C., Muddathir A., Adil H., Elhassan G., Ibrahim M., David-Polo-Orozco J., Cardone R., Reshkin S., Harguindey S. Glycolysis, tumor metabolism, cancer growth, and dissemination. A new pH-based etiopathogenic perspective and therapeutic approach to an old cancer question. Oncoscience, 2014;1(12):777–802. DOI: 10.18632/oncoscience.109.

19. Vander-Heiden M., Cantley L., Thompson C. Understanding the Warburg effect: the metabolic requirements of cell proliferation. Science 2009;324(5930):1029–1033. DOI: 10.1126/science.1160809.

20. Alfarouk K., Shayoub M., Muddathir A., Elhassan G., Bashir A. Evolution of Tumor Metabolism might Reflect Carcinogenesis as a Reverse Evolution process (Dismantling of Multicellularity). Cancers 2011;3(3):3002–3017; https://doi.org/10.3390/cancers3033002.

